# Utilizing occupancy-detection models with museum specimen data: promise and pitfalls

**DOI:** 10.1101/2021.12.05.471316

**Authors:** Vaughn Shirey, Rassim Khelifa, Leithen K. M’Gonigle, Laura Melissa Guzman

## Abstract

Historical museum records provide potentially useful data for identifying drivers of change in species occupancy. However, because museum records are typically obtained via many collection methods, methodological developments are needed in order to enable robust inferences. Occupancy-detection models, a relatively new and powerful suite of methods, are a potentially promising avenue because they can account for changes in collection effort through space and time. Here we present a methodological road-map for using occupancy models to analyze historical museum records. We use simulated data-sets to identify how and when patterns in data and/or modelling decisions can bias inference. We focus primarily on the consequences of contrasting methodological approaches for dealing with species’ ranges and inferring species’ non-detections in both space and time. We find that not all data-sets are suitable for occupancy-detection analysis but, under the right conditions (namely, data-sets that span long durations and contain a high fraction of community-wide collections, or collection events that focus on communities of organisms), models can accurately estimate trends. Finally, we present a case-study on eastern North American odonates where we calculate long-term trends of occupancy by using our most robust workflow.

## Introduction

Global change processes are contributing to the rapid restructuring of biodiversity across the planet (Araújo and Rahbek, 2006; Bellard et al., 2012). At the same time, data regarding species occurrence are becoming more widely available through the digitization of museum collections (Hedrick et al., 2020) and citizen science platforms (Dickinson et al., 2012). At the time of writing, two large databases of species occurrences, the Global Biodiversity Information Facility (GBIF) and Integrated Digitized Biocollections (iDigBio), contain over 1.5-billion records of species occurrences across the planet. These records may be rich in detailed information including date of collection, location, taxonomic determinations, and collector names. However, because museum data are aggregated records from many different collectors and time periods, they do not derive from a single sampling design and, further, the sampling design associated with each specimens is rarely noted. Consequently, they can contain numerous spatiotemporal and taxonomic biases (Isaac et al., 2014). For example, sampling of opportunistic (henceforth, ”unstructured”) data is often uneven in space and time and may be spatially concentrated around regions of high human population densities (Mair and Ruete, 2016; Tiago et al., 2017; Daru et al., 2018; Shirey et al., 2021). Additionally, charismatic taxa are often oversampled with respect to the diversity of their respective clades, leading to occurrence shortfalls, or fewer than expected occurrences given the diversity of a particular clade, in hyperdiverse groups such as arthropods (Troudet et al., 2017; Callaghan et al., 2021). In some groups, such as North American butterflies, undersampling is prolific in regions that are forecasted to experience the most dramatic changes in climate (Shirey et al., 2021). Given these biases, proper treatment of unstructured data can lead to misleading inferences (Guzman et al., 2021; Larsen and Shirey, 2021). Thus, it is imperative that we develop statistical frameworks for analyzing unstructured data in order to afford researchers the ability to test hypotheses related to how global change is impacting species over long time-periods and large spatial extents.

Species observation is rarely perfect so, even when a given species is present at a site, the observer may fail to detect it during any given survey (e.g., animals with cryptic phenotype or behavior may be difficult to see). Occupancy-detection models are a powerful statistical framework for disentangling this imperfect processes of observation from actual species’ occurrence or ”occupancy” (MacKenzie et al., 2003) and these models have been widely used for species’ distribution modeling in ecology (Kéry and Schaub, 2011). Occupancy models have been extended to model entire communities (so called, ”multi-species occupancy models”) and to model one or more species across multiple seasons (so called, ”dynamic occupancy models”) (Kéry and Schaub, 2011). These models have improved our ability to explore spatial and temporal patterns of biodiversity and aided in identification of drivers of global change (Kéry and Royle, 2020).

Occupancy models are potentially well-suited to handle the inherent biases in species observations present in unstructured data (Erickson and Smith, 2021). A major challenge when doing so is accounting for the fact that records of when and where historical collectors sampled are rarely available. Collections that did not yield specimens, or did but the specimens were never archived in a museum, are typically unreported, because no physical specimens exist in the museum collection. This shortcoming is further exacerbated by variable (and also unknown) individual collector behaviors. For example, some collectors focus on particular taxonomic groups while others collect entire communities (de Siracusa et al., 2020; Di Cecco et al., 2021). Species’ traits may also influence the probability of detection or ”allure” and, thus, collection outcomes (Johnston et al., 2014). In all, physical specimens and their metadata represent a fragmented history of potential collection events that researchers must carefully piece together to reconstruct historical patterns.

Researchers have developed some best-practices for overcoming some of these challenges for presence-only data. For instance, it might be appropriate to infer non-detections of one (or more) species at a given site based on known detections of other species at that same site (Kéry, 2010; van Strien et al., 2013a; Kamp et al., 2016a). Constraining detection-non-detection records only to those sites that are plausibly within the range of a species’ geographic distribution has also been shown to improve model performance (Guzman et al., 2021). Combined, these steps not only improve estimates of occupancy and detection, but can also reduce the computation time of analyses. However, we lack clear guidelines that highlight best practices for avoiding the potential biases introduced by variable observer behavior, temporal trends in visitation, and species distributions and traits.

Here, we use a simulation framework to explore the performance of occupancy-detection models when applied to data that contain many of the patterns likely present in natural history collections data. Specifically, we explore how changes in collector visit frequency, species’ occupancy, and species’ detection impact model performance for data-sets containing presence only records across a community of species, each with its own geographic range. We consider the consequences of 1) incorporating estimated species’ ranges and 2) inferring non-detections of some species from recorded detections of others on estimated trends in occupancy and detection. We synthesize our findings by providing guidelines for steps that could be taken to reduce bias when applying similar occupancy-detection models to empirical data-sets. Finally, we apply our scenarios to a real-world data-set containing detection records for eastern North American odonates (dragonflies and damselflies).

## Methods

### Data simulation

We simulate a community of *N* species, each inhabiting some or all of *J* potential sites over *K* time periods, which we call ”eras.” Eras could correspond to single years or could span multiple years. We suppose that each era is further broken into *I* time intervals, during which collectors may visit sites to sample some or all of the species present (commonly used terms in this manuscript are presented in the glossary of terms Box 1).

#### Table 1: Box: Glossary of terms commonly used throughout this manuscript.

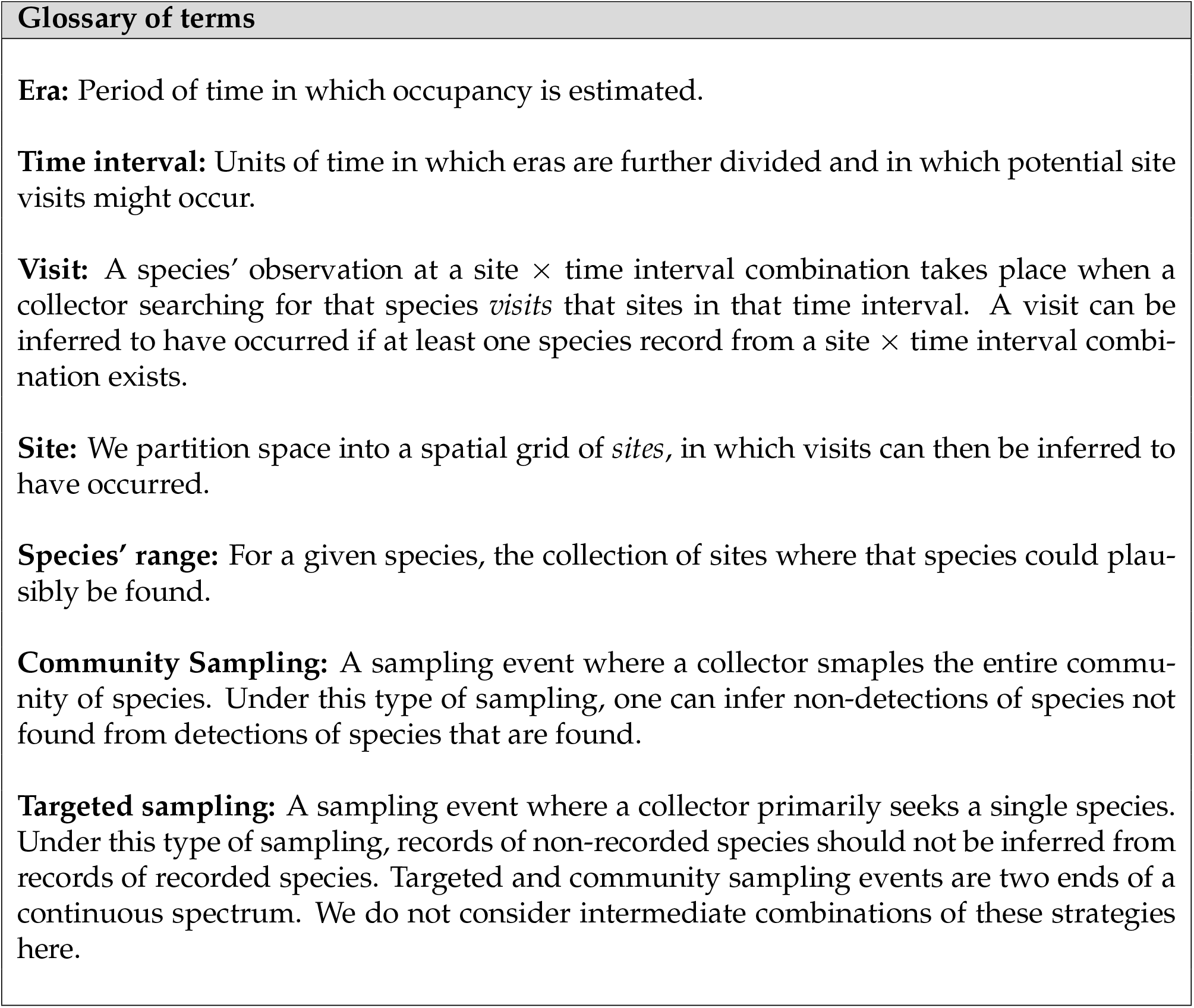

Because species often differ in their spatial extent, in some of our scenarios, we simulate a unique ”geographic” range for each species. Specifically, species *i* potentially occupies sites *R_i_* = {*s*_*i*,1_,*s*_*i*,2_, ⋯, *s_i,r_i__*}, where *r_i_* ∈ {1,2, ⋯, *J*}. We refer to *R_i_* as the ”range” of species *i*. To simulate each species range, we generated polygons where size was determined using a Poisson distribution. Ranges were constructed by placing a series of random vertices on a defined grid and by drawing a concave polygon around these vertices. The size of the range was defined by a scaling factor, *r_i_* which was drawn from a Poisson distribution as:

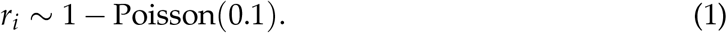

For site *j* within the range of species *i*, we simulate occupancy in era *k* by drawing from a Bernoulli distribution with success probability *ψ_i,j,k_*, where

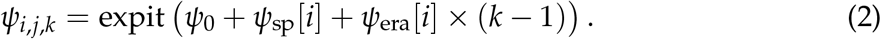

Here *ψ*_0_ denotes the baseline occupancy (on the linear scale), *ψ*_sp_[*i*] is a random species-specific intercept, and *ψ*_era_[*i*] is a random species-specific effect of era (a positive value of *ψ*_era_[*i*] would indicate that species *i* is increasing in occupancy through time). We assume that *ψ*_sp_[*i*] and *ψ*_era_[*i*] are normally distributed, such that:

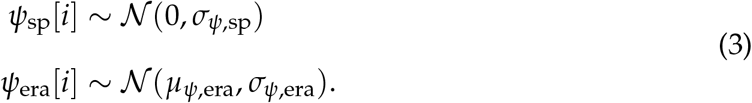

Here, *σ*_*ψ*,sp_ denotes the interspecific variability in occupancy, *μ*_*ψ*,era_ denotes the mean effect of era on species occupancy, and *σ*_*ψ*,era_ denotes the variability in species’ temporal occupancy trends.

Because sampling effort can change through time, we assume that the probability that a site is visited in a time interval can change across eras. Further, because some collectors may record all species that they detect, while others may record only a single species, all site visits are not equivalent. If a collector records all species (we call this ”community sampling”), then a visit to a site in a given time interval is a relevant visit *for all species*. However, if a collector only records a single species, ignoring others (we call this ”targeted sampling”), then a visit to a site in a given time interval is only a meaningful visit for the recorded species. In our data simulation, we suppose that, in era *k*, site *j* receives a ”community sampling” visit in each time interval with probability

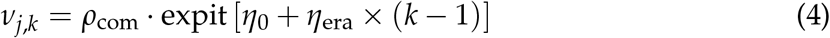

and when such a visit occurs, each species is recorded as either detected or not detected. Here, *η*_0_ denotes the baseline probability of a community sampling visit (on the linear scale) and *η_era_* is the effect of era on visitation probability. This latter term allows the probability of site visitation to change systematically across eras. *ρ*_com_ lets us tune the fraction of community sampling visits relative to targeted sampling events; if *ρ*_com_ = 1, all visits are community sampling events, whereas, if *ρ*_com_ = 0, all sampling events are targeted sampling events. By parameterizing *ρ*_com_ as a function of model parameters, one could also allow collector behaviour to change across space or through time, however, that is beyond our scope here.

Only time intervals (at each site) that do not receive a community sampling visit can be subjected to targeted sampling visits (a targeted sampling visit will not add any additional information if a community sampling visit also occurs in the same time interval). Targeted sampling visits are, by their nature, species-specific. The probability that, in era *k*, site *j* receives a targeted sampling visit for species *i* is given by

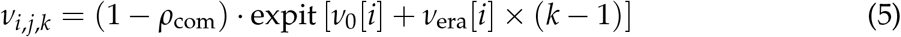

where *ν*_0_[*i*] denotes the baseline probability of a targeted sampling visit searching for species *i* (on the linear scale) and *ν*_era_[*i*] denotes the effect of era on the probability of targeted sampling visits searching for species *i*. We assume that both *ν*_sp_[*i*] and *ν*_era_[*i*] are normally distributed, such that

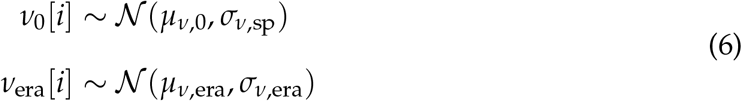

where *μ*_*ν*,0_ denotes the mean probability, across species, of a targeted sampling visit (in the first era), *μ*_*ν*,era_ denotes the mean effect, also across species, of era on the probability of targeted sampling visits, and *σ*_*ν*,sp_ and *σ*_*nu*,era_ denote the interspecific variation in visitation probability and variation in temporal change in visitation, respectively. In our simulation, we let *η*_0_ equal *μ*_*ν*,0_, and *η*_era_ equal *μ*_*ν*,era_ so that both the probability of community and targeted sampling events increase or decrease simultaneously, but this does not need to be the case. For example, the probability of community sampling events could increase across eras, while the probability of targeted sampling events can go down. Exploring that scenario could be interesting but beyond the scope of this manuscript. Because *η*_era_ is equal to *μ*_*ν*,era_, variations of *μ*_*ν*,era_ (and therefore *η*_era_) will simply be labelled as *μ*_*ν*,era_.

Finally, for each species, we simulate detection during the time intervals where visits occurred. For site site *j* ∈ *R_i_*, we draw species’ *i* detections across the visits that occurred and could have detected that species (e.g., community sampling visits or targetting sampling visits aimed at species *i*) in era *k* from a Bernoulli distribution with probability *p_i,j,k_* where

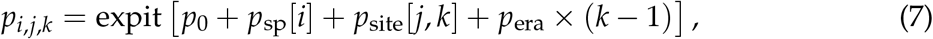

where *p*_0_ is the baseline detection probability (on the linear scale), *p*_sp_ [*i*] is a random species-specific intercept, *p*_site_[ *j*, *k*] is a random site-specific intercept that varies by era, and *p*_era_ is an overall effect of era. *p*_site_ [*j, k*] allows spatiotemporal variability in detection probability (changing across sites and eras), and helps account for the variation that is inherent in sample effort across space and time in opportunistic historical data-sets. *p*_era_ allows detection to change systematically through time, as has likely occurred in many groups as sampling techniques have improved, for example. We assume that both *p*_sp_ and *p*_site_ are normally distributed, such that:

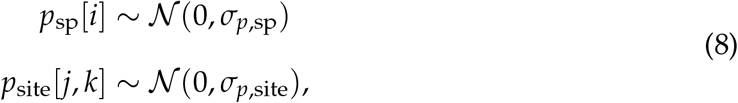

where *σ*_*p*,sp_ denotes the interspecific variation in detection and *σ*_*p*,site_ the spatiotemporal (site and era) variation in detection.

We considered (i) three different numbers of eras (*K*), (ii) five different fractions of community sampling events (*ρ*_com_), (iii) three different trends in visitation probability (*μ*_*ν*,era_; increasing, decreasing, and stable), and (iv) identical versus variable species ranges for a total of 135 simulation scenarios (specific parameter values are shown in Table S1 and Fig. 1). We conducted all data simulations in R (code is available at https://github.com/lmguzman/occ_historical).

**Figure 1:**
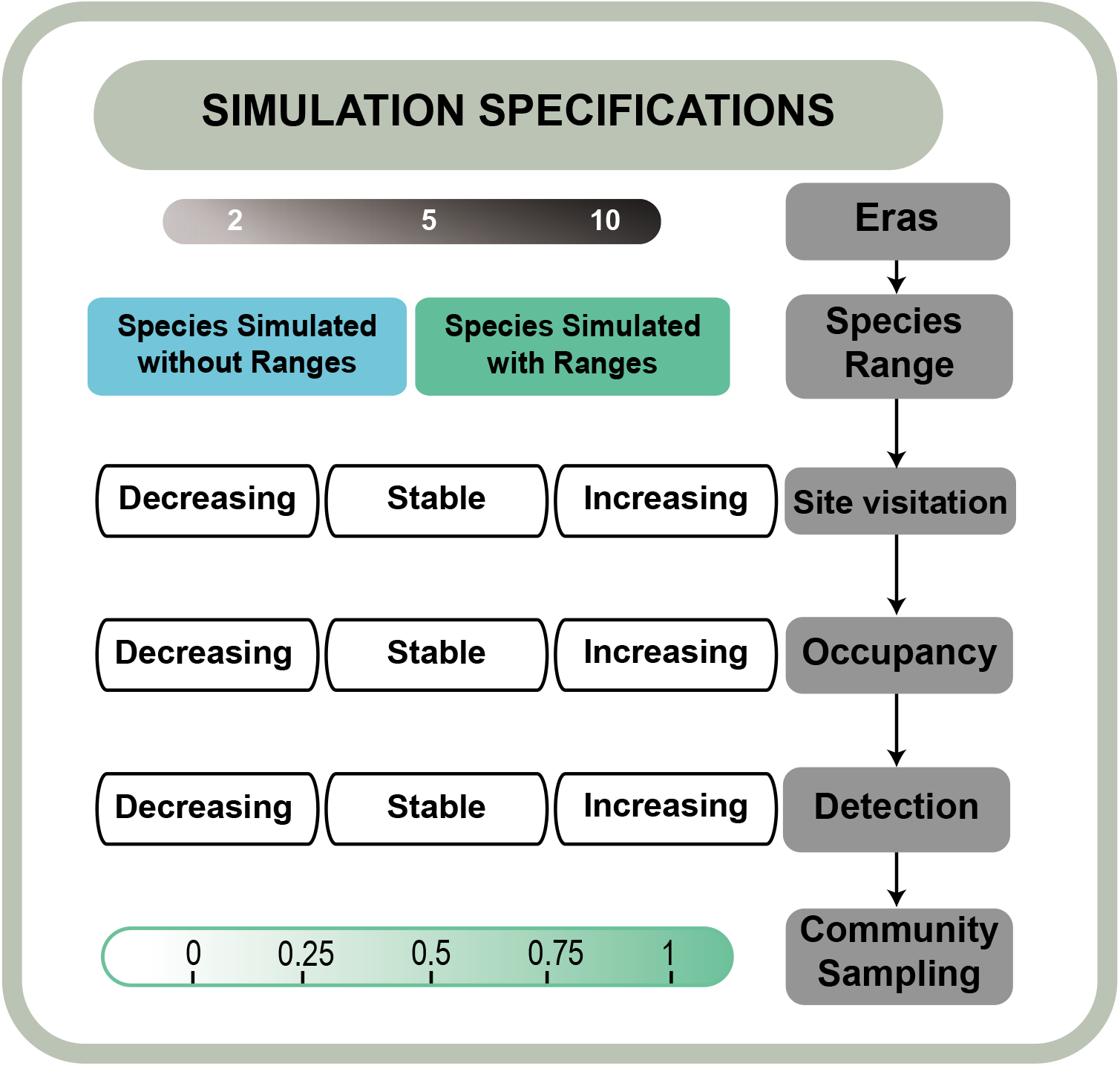
Schematic depicting data simulation decisions. Visit, occupancy, and detection probabilities all have three levels: decreasing, stable, and increasing with time (−0.1, 0, 0.1). The probability that visits are community sampling events is: 0, 0.25, 0.50, 0.75, 1. All five parameters were varied sequentially, which resulted in 270 multi-species detection-non-detection data-sets (N=2*3*3*3*5). We also simulated data with two, five, and 10 eras.

### Occupancy-detection models

We built an occupancy model that reflect the structure of the simulated data-sets, except that we do not incorporate site visitation history or community sampling information, as this information is typically not easily accessible (or accessible at all) for empirical data-sets. We provide a detailed descriptions of our occupancy models in the Supplementary Information.

Using our simulated data-sets, we constrast the performance of six different decision workflows one might follow when applying an occupancy model to a historical data-set. We provide an overview of these below and they are also shown graphically in Fig. 2.

- *WF*_all,all_: we model all sites for all species across all time intervals, even when some of those time intervals did not contain any known species detections.
- *WF*_all,detected_: we model all sites for all species, but each site is only modelled over the time intervals where at least 1 species was detected. Here, we infer non-detections of non-observed species when at least one other species was detected at a site in a given visit. This is a fair assumption when most collectors sample the entire community (*ρ*_com_ = 1), but not when they mostly target individual species (*ρ*_com_ = 0).
- *WF*_all,visits_: we model all sites for all species, but each site only over the time intervals where we know visits occurred (i.e., we model the true site visitation history which we are only able to do because we are working with simulated data). This case provides a benchmark for a ’best information’ workflow.
- *WF*_range,all_: we model each species over only the sites that fall within its range and each site is modelled across all time intervals (this is an analog to *WF*_all,all_ above, but where species’ ranges are estimated and then incorporated).
- *WF*_range,detected_: we model each species over only the sites that fall within its inferred range and, for each of those sites, only over time intervals where at least 1 species was detected (this is an analog to *WF*_all,detected_ above, but where species’ ranges are estimated and then incorporated).
- *WF*_range,visits_: we model each species over only the sites that fall within its inferred range, but each site only over the time intervals where we know visits occurred (this is an analog to *WF*_all,visits_ above, but where species’ ranges are estimated and then incorporated).

**Figure 2:**
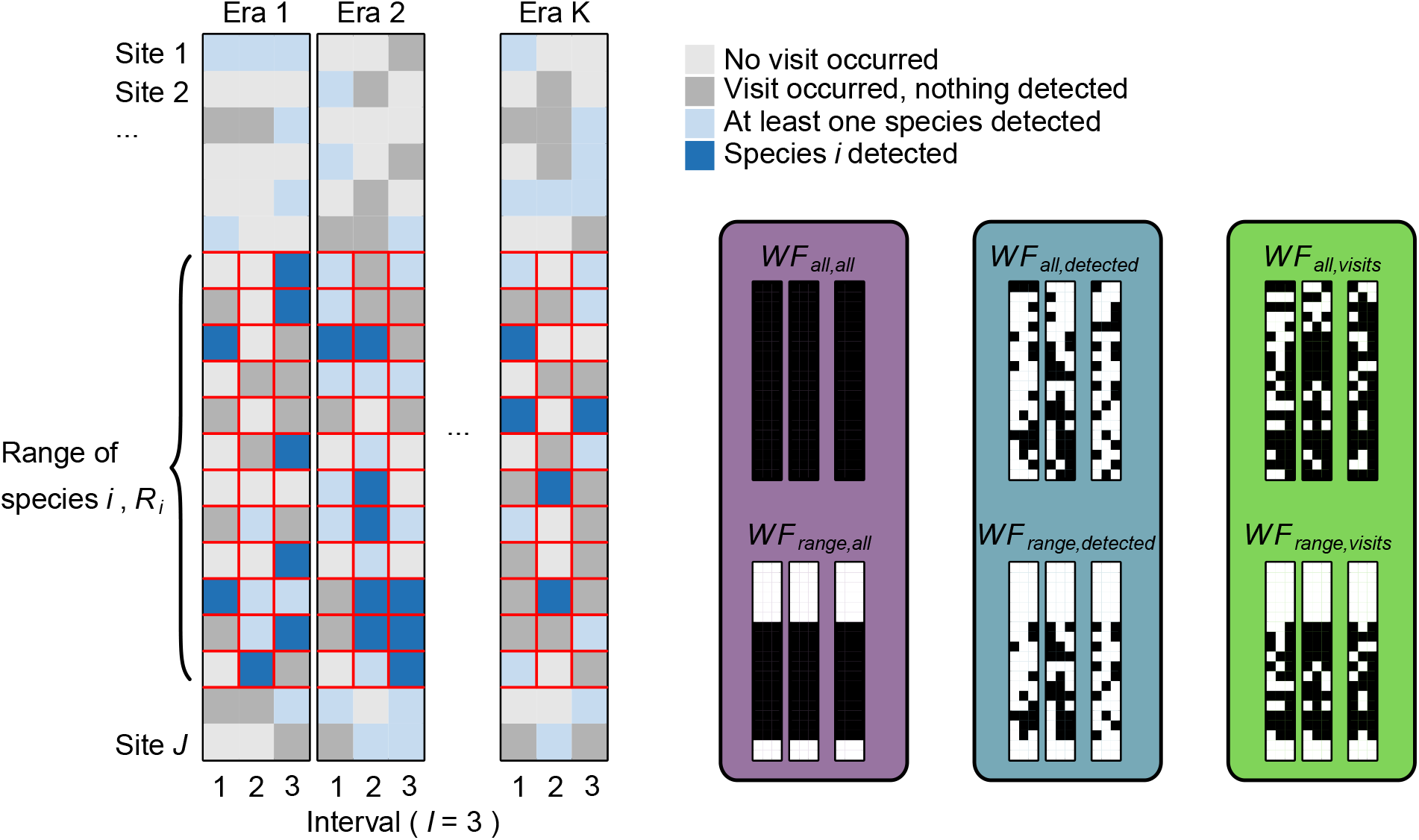
Schematic depicting data processing workflows. Workflows differ in which sites and time intervals they model. Small black and white panels show, for species *i*, which site × time interval combinations from the larger panels are modelled in each workflow, with black denoting a modelled site × time interval combination and white a non-modelled combination. For example, *WF*_range,all_ models all time intervals for every site within the range of species *i*, but none of the sites outside of it. Colours in large boxes behind small panels correspond to colours in subsequent figures.

We compared these different processing workflows by assessing accuracy of model-estimated parameter values against their true values using root mean squared error (RMSE) across five replicates.

### Case Study: Eastern North American Odonates

#### Study organism

Odonates are globally widespread aquatic insects that live in a wide range of freshwater systems. They are charismatic, easy to detect and identify, and popular among nature enthusiasts. Both larvae and adults are generalist predators and interact with a large number of aquatic and terrestrial taxa. In North America, there are 462 species (136 Zygoptera and 326 Anisoptera), with 336 species on the east coast (Paulson, 2011). Odonates are sensitive to environmental change, particularly climate warming, and evidence suggests that they respond by shifting their phenology and geographic distribution (Hickling et al., 2005; Hassall et al., 2007), making them interesting sentinels of climate change.

#### Odonate data-set

We selected 27 states in Eastern North America from GBIF (https://doi.org/10.15468/dl.cabqrc). The data comprise 152,863 records of 368 species observed during 1970 – 2020 and show a clear increase in the number of records through time, with 70.3% of the records occurring in the last two decades (2000 – 2020). We partitioned records into ”sites” by overlaying a grid of 2° × 2° cells and filtered to include only species with more than 100 occurrences (194 species) across 90 sites and five eras (decades), each further partitioned into five two year time intervals.

#### Analysis of the odonate data-set

We provide guidelines for researchers when analysing historical museum records with occupancy models (Box 2) and we demonstrate application of these steps using the above-described Odonate data-set. Specifically, we use a multispecies occupancy model, as described in the supplementary materials. For each species, we construct the geographic range by drawing a convex polygon around all occurrences records. We process the data under the *WF*_range,detected_ (best case scenario on simulated data, excluding *WF*_range,visits_ which is not possible for empirical data), *WF*_all,detected_ which ignores underlying species ranges, and *WF*_range,all_ which models all visits, regardless of whether or not visits likely occurred. We also assessed how many of the records in our data-set likely originated via community sampling vs. targeted sampling visits. To do this, we grouped records collected on the same day and within 1km of one other. We assumed that a community sampling event occurred if more than one species was present in each grouping. While this is likely a rough approximation of the actual fraction of community sampling events, it lets us quickly approximate this quantity without having to use more time-consuming methods like a detailed analysis of collector identities, etc.

##### Table 2: Box: Steps that researchers can take to analyse historical record data using occupancy models, illustrated using a data-set of Odonate occurrences.

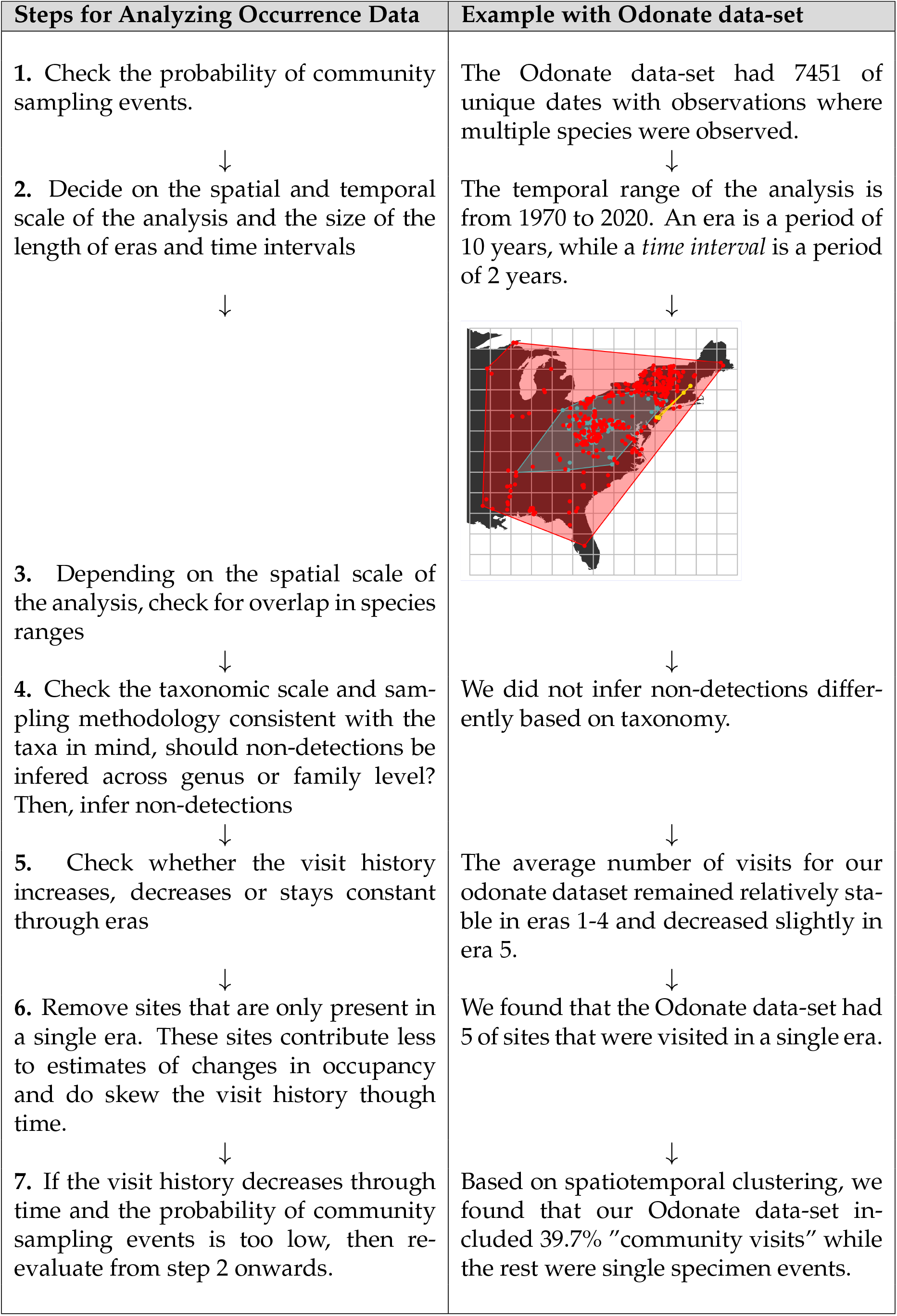

| Steps for Analyzing Occurrence Data | Example with Odonate data-set |
| --- | --- |
| 1. Check the probability of community sampling events. | The Odonate data-set had 7451 of unique dates with observations where multiple species were observed. |
| ↓ | ↓ |
| 2. Decide on the spatial and temporal scale of the analysis and the size of the length of eras and time intervals | The temporal range of the analysis is from 1970 to 2020. An era is a period of 10 years, while a *time interval* is a period of 2 years. |
| ↓ | ↓ |
| 3. Depending on the spatial scale of the analysis, check for overlap in species ranges |  |
| ↓ | ↓ |
| 4. Check the taxonomic scale and sampling methodology consistent with the taxa in mind, should non-detections be inferred across genus or family level? Then, infer non-detections | We did not infer non-detections differently based on taxonomy. |
| ↓ | ↓ |
| 5. Check whether the visit history increases, decreases or stays constant through eras | The average number of visits for our odonate dataset remained relatively stable in eras 1-4 and decreased slightly in era 5. |
| ↓ | ↓ |
| 6. Remove sites that are only present in a single era. These sites contribute less to estimates of changes in occupancy and do skew the visit history though time. | We found that the Odonate data-set had 5 of sites that were visited in a single era. |
| ↓ | ↓ |
| 7. If the visit history decreases through time and the probability of community sampling events is too low, then re-evaluate from step 2 onwards. | Based on spatiotemporal clustering, we found that our Odonate data-set included 39.7% "community visits" while the rest were single specimen events. |

We performed all analyses with R4.1.1 (R Core Team, 2019). We used the nimble package to compute our models (de Valpine et al., 2017). All of the code utilized in our study is available at https://github.com/lmguzman/occ_historical.

## Results

### Simulation Study

The choice of data processing workflow directly affects inferences made by occupancy models. Not surprisingly, workflows that only include site × time interval combinations where visits actually occurred (*WF*_all,visits_ and *WF*_range,visits_) outperformed models that inferred site visitation and this was invariant to the fraction of visits that represent community sampling events (Fig. 3). The performance of workflows that infer species’ non-detections by either modelling all time intervals (*WF*_all,detected_) or only time intervals where at least one species was detected (*WF*_all,all_) depends on the fraction of visits that are community sampling events. Specifically, when all site visits correspond to community sampling events (*ρ*_com_ = 1), inferring species non-detections based on positive detection of other species is a sensible action and, consequently, *WF*_all,detected_ performs as well as *WF*_all,visits_ (Fig. 3c,d,e,f). In contrast, when all site visits are targeted sampling events, (*ρ*_com_ = 0), detections of one species provide little information about non-detections of other species and, consequently, both *WF*_all,detected_ and *WF*_all,all_ yield biased estimates of occupancy change through time (purple and blue curves are much higher than green curve for small values of *ρ*_com_ in Fig. 3).

**Figure 3:**
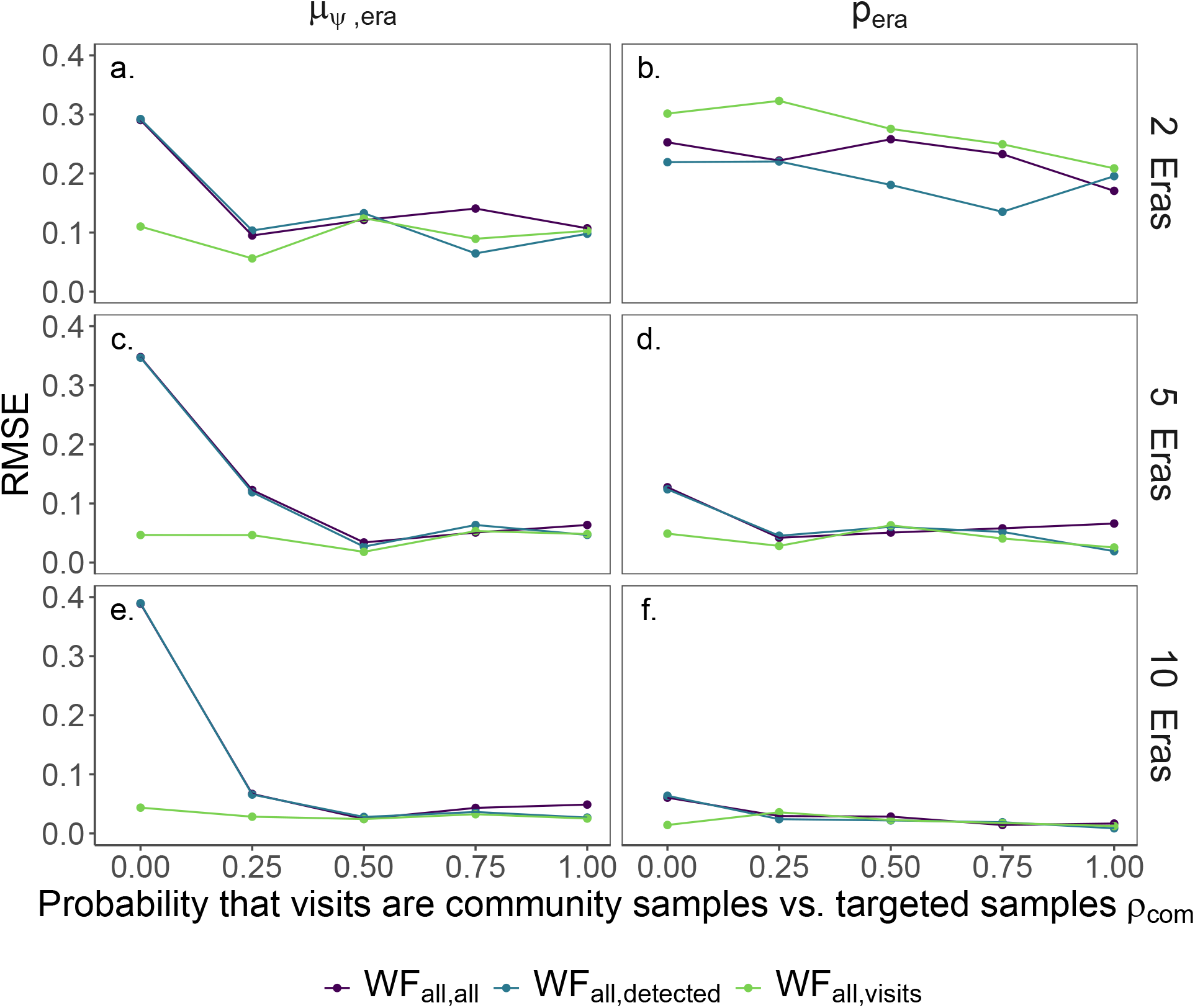
When we only model time intervals where visits actually occurred (*WF*_all,visits_, green line), estimates of change in occupancy (*μ*_*ψ*,era_) and detection (*p*_era_) are generally more accurate (lower root mean square error RMSE) compared to workflows where we either model only visits with at least one species detected (*WF*_all,detected_, blue line) or where we model all time intervals (*WF*_all,all_, green line). This difference is largest when only the fraction of the visits that were community sampling events is low (small *ρ*_com_). Across the board, models that span a greater number of eras generally yield more precise estimates. *μ*_*ν*,era_ = 0.

We found that model error increased as the minimum fraction of visits that correspond to community sampling events decreased and, further, that the extent of this error depended on (i) the number of eras, (ii) whether species’ ranges were incorporated into the analysis, (iii) the extent to which site visitation probability changed through time, and (iv) the extent to which occupancy and/or detection probability changed through time. We will discuss these processes next.

In general, estimated temporal changes in occupancy and/or detection probability through time are more accurate when data comprise a greater number of eras (larger K) and this conclusion is true across data processing workflows (Fig. 3). Model estimates for data-sets spanning ten eras were able to better estimate changes in occupancy and detection through time when community visits comprise a smaller fraction of the total site visits (lower *ρ*_com_) than estimates for data-sets spanning five eras (Fig. 3c,d,e,f). Estimates for changes in detection through time were, however, less sensitive to both the fraction of visits that comprise community sampling events and to the workflow (Fig. 3, Fig. 4).

**Figure 4:**
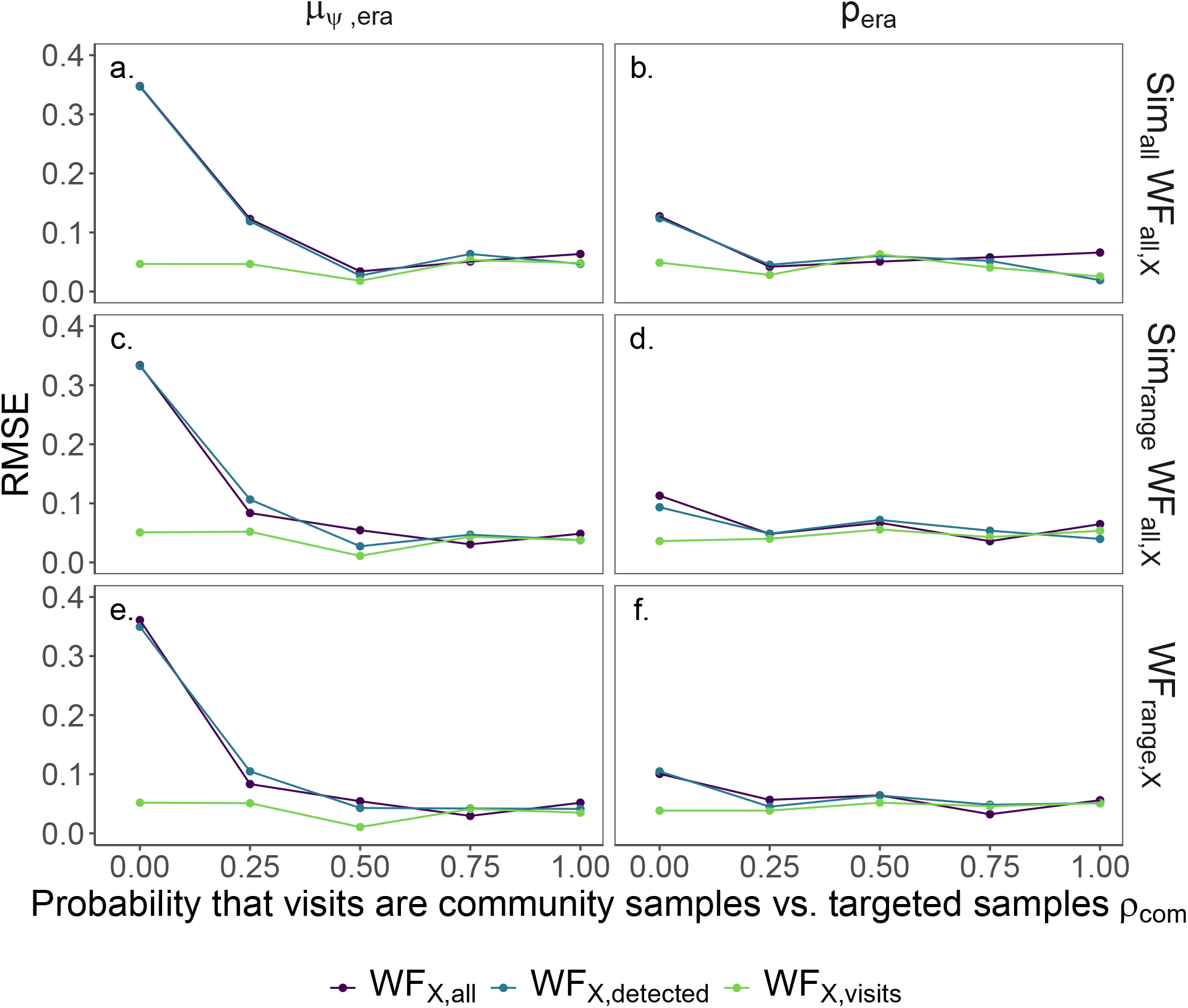
Changes in occupancy (*μ*_*ψ*,era_) and detection (*p*_era_) through time have less error (root mean squared error RMSE) as the probability of community visits *ρ*_com_ increases. The coloured lines are the workflows regarding to the visitation history where X just stands in for any range workflow. The rows show the effect of the workflows with regards to the range and the underlying simulation. Sim_all_ is when all of the species were simulated to plausibly occupy all sites, while Sim_range_ is when species were simulated to have ranges. *WF*_all,X_ is the data processing workflow where all sites are included in the occupancy model and *WF*_range,X_ is the data processing workflow where the only sites that are included in the model are the ones that fall within a species range. *μ*_*ν*,era_ is zero, Number of eras is five.

We simulated data-sets either by assuming that every species could potentially occupy every site or by assuming species were constrained to different (possibly overlapping) geographic ranges. We subsequently differentiated our data processing workflows into those that ignore potential differences in species ranges (*WF*_all,all_, *WF*_all,detected_, and *WF*_all,visits_, which we collectively label as *WF*_all,X_) and those where each species is modelled over only the sites in its range (*WF*_range,all_, *WF*_range,detected_, and *WF*_range,visits_, which we collectively label as *WF*_range,X_). When all species could plausibly occupy all sites, *WF*_all,X_ and *WF*_range,X_ become equivalent. We found that workflows that consider underlying species’ ranges (either *WF*_all,X_ when all the species were simulated to be plausibly occupy all sites or *WF*_range,X_ when species ranges were simulated) generally had lower error rates in changes of occupancy through time than the case where the data processing workflow mismatched the underlying species ranges (using *WF*_all,X_ when species ranges were simulated) (Fig. 4a,c).

When the probability of site visitation decreased through time, we found that workflows that attempted to infer and model species’ detections based on recorded presences of other species (e.g., *WF*_range,detected_) yielded better occupancy and detection estimates than workflows that simply modeled all species across time intervals (e.g., *WF*_range,all_). This pattern is most obvious when most visits correspond to community sampling events (high *ρ*_com_, Fig. 5, top row). This effect disappeared when site visitation probability remained constant or increased through time (Fig. 5 bottom rows). Generally, an increasing probability of site visitation through time improves model inferences compared to decreasing probability of site visitation when a lower fraction of the visits correspond to community sampling events. For instance, models yield accurate estimates of occupancy and detection when only 25% of site visits are community sampling events, as long as the site visitation probability increased through time, the number of eras was high (*K* = 10), the data processing workflow took into account the underlying species ranges, and species’ non-detections were only inferred when other other species were detected (i.e., *WF*_range,detected_).

**Figure 5:**
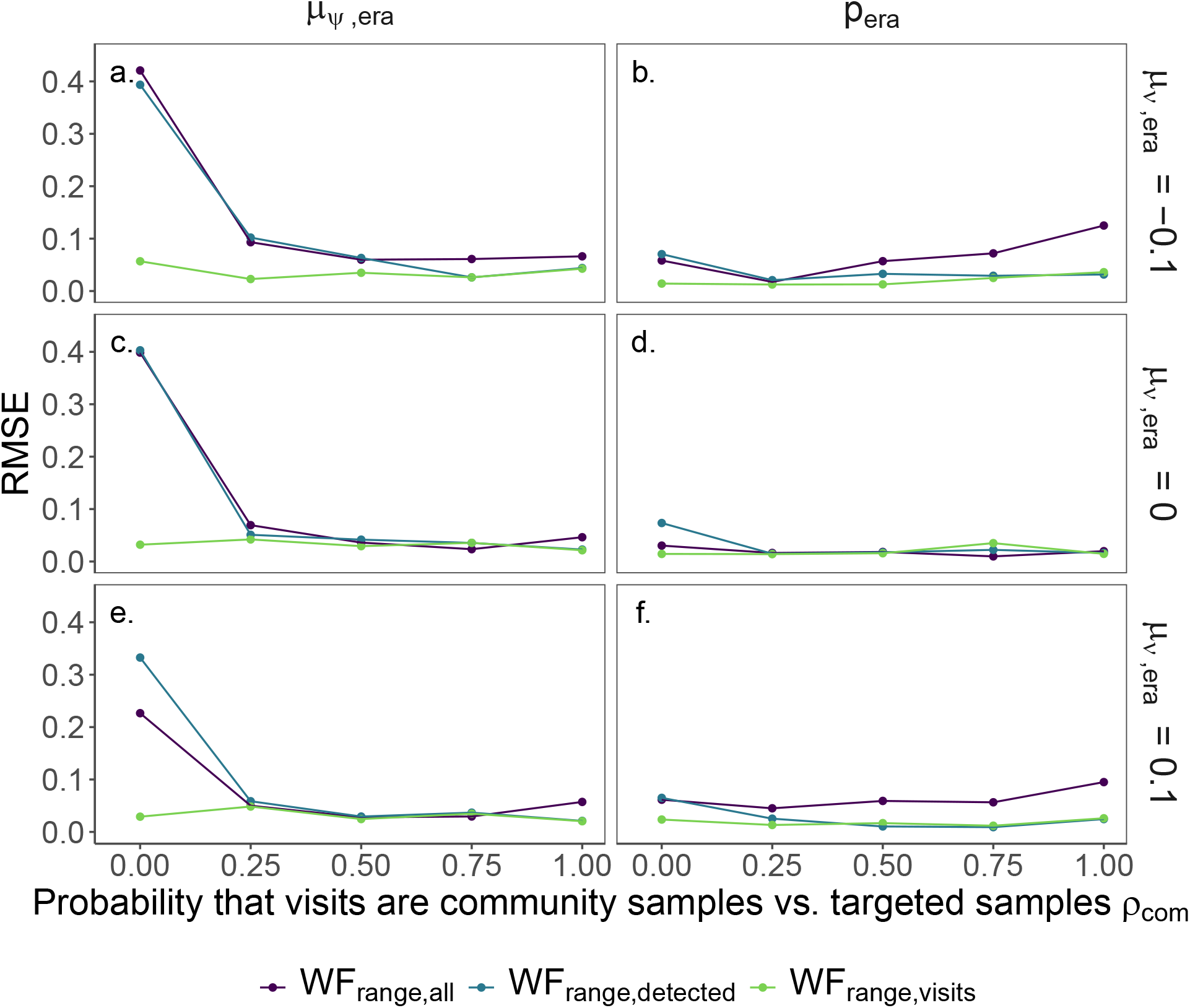
Changes in occupancy (*μ*_*ψ*,era_) and detection (*p*_era_) through time have less error (root mean squared error RMSE) using the workflow *WF*_range,visits_ than *WF*_range,detected_ and *WF*_range,all_. This error depends on the probability of community visits *ρ*_com_ and the change in visit history through time (*μ*_*ν*,era_). Number of eras is ten.

Because real-world changes in quantities like site visitation rate, occupancy, detection, and collector behavior are potentially changing through time, we ran a full suite of occupancy models that included 3,750 workflow-simulation combinations with five replicates per combination. Root-mean squared errors (RMSEs) from this full set of workflow-simulation configurations revealed that overestimation of occupancy declines are likely to occur when the fraction of visits that are community samples is low (0 – 0.25) and this is especially true when detection and visit probability decrease through time (Supplemental Figures S1, S2, and S3).

### Case Study: Eastern North American Odonates

To examine occupancy trends for eastern North American odonates from the 1970s through 2010s, we ran occupancy models on data prepared using each of the processing workflows WFrange,detected, *WF*_all,detected_, and *WF*_range,all_. We found that the proportion of site × era × time interval combinations that were likely community samples in this data was ~ 39.7%. Because our conclusions from simulated data suggest that this might be too low for models to generate non-biased estimates, we re-filtered our data, applying stricter criteria. Specifically, we restricted our records to only those from site × era × time interval combinations where at least 50% of sampling events were inferred to be community sampling events (i.e., multiple species detected on the same day within 1km × 1km from one other). This led to an estimated global proportion of community samples equal to 76.6%. We then re-ran an occupancy model using our best performing workflow (*WF*_range,detected_) on this restricted data-set.

In general, we found that mean occupancy increased over the past 4 decades. However, species trends were highly variable, with many increasing while others decreased. By using a range-restricted workflow, our odonate results became less extreme in terms of overall, species-specific occupancy change from the 1970s to 2010s (Fig. 6). For example, using an *WF*_all,detected_ workflow yields 65 species that are either increasing or decreasing by at least 25-percent. Restricting our workflow to *WF*_range,detected_ reduces this figure to 22 species (24 species for *WF*_range,detected_ with site/era/visit combinations that were clustered, community visits - 6d).

**Figure 6:**
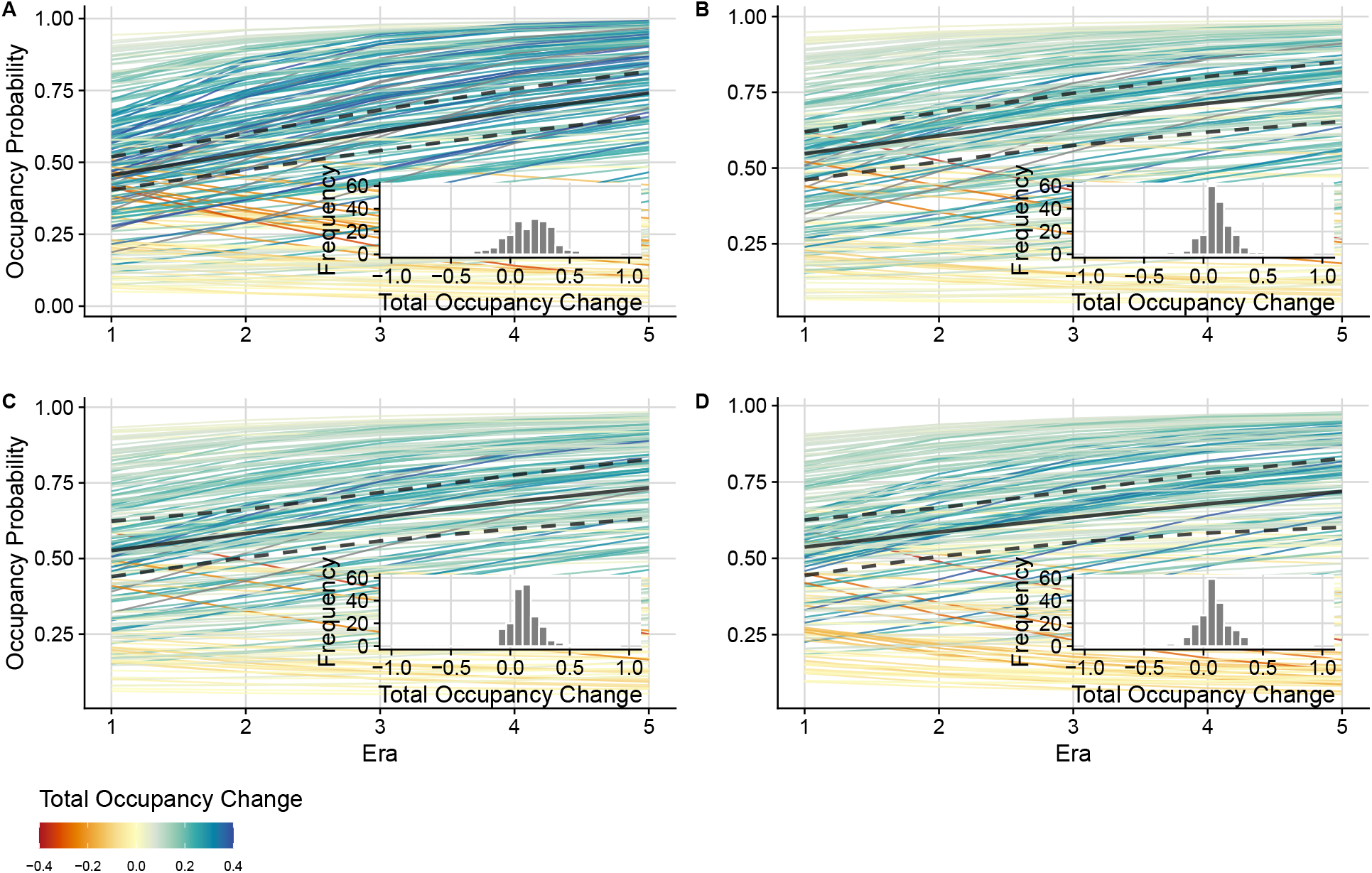
Species-specific (colored lines) and community (black line) trends for eastern North American odonates under three data processing workflows: (a) *WF*_all,detected_, (b) *WF*_range,all_, (c-d) *WF*_range,detected_. In (d), we further restrict analyses to only site × era × time interval combinations where the proportion of clustered collection events containing 2 or more species exceeds 0.5. Lines are coloured to indicate occupancy trend through time (difference from Era 1 to Era 5). Inset histograms show these occupancy differences. Black dashed lines denote 95% Bayesian Credible Intervals.

## Discussion

We assessed the performance of different data-processing workflows when applying multi-species occupancy models to simulated data-sets in order to identify a set of best practices guidelines one might follow when analyzing historic museum records or opportunistic data. Our work here confirms the importance of restricting species-specific analyses across large geographic scales to those species’ ranges. Further, inferring non-detections of species from other species’ detections emerged as a better approach than simply modelling all possible time intervals at all sites. Our analysis also determines the extent to which patterns in data sampling, such as changing collector behaviour, can undermine our ability to infer trends using occupancy models from historic records. In addition, workflows that bin occurrence data into fewer eras are generally less accurate.

Inferring non-detections of some species based on recorded observations of other species is becoming a more frequent practice when working with presence-only occurrence data, but specific approaches differ. For example, van Strien et al. (2013b); Kamp et al. (2016b); Powney et al. (2019) inferred non-detections for a particular species if a different species was observed at that same site on the same date. Outhwaite et al. (2019) used the same approach, but only inferred non-detections for species from the same taxonomic group as the observed individual(s). Analyses using these approaches have mostly taken place in Europe and with taxonomic groups where community science approaches yield structured data sets, for example those from regional monitoring schemes (van Swaay et al., 2008). For example, Johnston et al. (2021) found that well curated checklists combined with incomplete checklists were sufficient to enable reasonable estimates of occupancy, but not encounter rates. Here we have quantified the extent to which a data-set can comprise incomplete checklists or simply individual observations while still producing good parameter estimates. We found that, for the cases considered here, our workflows produced reliable estimates of occupancy trends through time when a least 50% of the samples derived from community sampling events.

Obtaining an approximate estimate of how frequently collectors sampled the entire community, versus only collecting or recording individual species, is a necessary step in assessing whether the methods we use here are likely to be appropriate. Unfortunately, direct measurement of this quantity is often impossible for large historic data-sets and, thus, we suggest that researchers approximate this metric to the best of their ability. In our Odonate analysis, we grouped records in space and time and assumed that specimens collected on the same day and nearby to one another were part of the same collection event. Based on our simulated data-sets, we concluded that reliable estimates for data-sets that span only two time periods require that greater than 50% of site visits must comprise community sampling events. In workflows that span a greater number of eras, this fraction could conceivably be as low as 25%, however, this likely depends on other data-specific trends and parameter estimates, and we would suggest that authors conducting such analyses always use simulation to guide their analyses. Our examination of interactions between changes in occupancy and detection probability over time as well as changes in visitation probability revealed that when detection probability or when visit probability decrease over time, overestimation of occupancy declines will likely occur in data-sets where the probability of community visits is less than 0.5 (Supplemental Figures S1, S2, and S3).

In applying our different workflows to a large historic data-set for eastern North American odonates, we demonstrate that neglecting to incorporate estimates of species’ ranges demonstrates how different workflow choices can impact downstream results with a real-world data-set. In particular, the results from this analysis show how data filtering can reduce the estimation of dramatic occupancy increases and declines (Fig. 6a vs. Fig. 6b,c,d). Overestimation of occupancy change is a known behavior of occupancy-detection models when heterogeneous observer effort is not accounted for (Kéry and Royle, 2020). Our findings generalize this conclusion to show that similar overestimation is possible when species are modeled outside of their ranges (using the Odonate example), when species are modelled in all time intervals, or when there is too few community sampling events (using simulated data).

Using the best workflow to analyze the Odonate dataset, we found that the majority of species had range expansions and a low number of species had range contractions in the eastern United States. This is mostly concordant with the assessment of the conservation status and population trends of the IUCN Red List at the global scale which lists all studied species as Least Concern with stable or increasing population trends. Odonates have a high ability to colonize new suitable habitats following the warming of cool areas (Hickling et al., 2005) or the creation of artificial sites (e.g. ponds for irrigation) (Simaika et al., 2016). The observed low range contractions might be due to their ability to rapidly recolonize areas where they were temporarily extirpated (Shiffer and White, 2014), which is often observed at the local scale in the studied area (Shiffer et al., 2014, 2015). Studies using occupancy models to assess the temporal pattern of the geographic distribution of odonates at a large scale are rare (Rocha-Ortega et al., 2020), however they may be possible under the occupancy-detection framework.

While we only use ”single-season” occupancy models in our analyses here, we also explored (but do not present) dynamic occupancy models which link each time periods (or eras, in our terminology) via extinction-colonization dynamics (Royle and Kéry, 2007). In general, we found that these models performed poorly unless the probability of community visitation was high. Missing visits/eras in the dynamic framework created multiple possibilities for parameter estimation, where a single era with missing data could represent extinction followed by colonization, or persistence; multiple consecutive eras with missing visits simply compounds the number of possibilities. We expect that many historical data-sets likely do not have enough data to make dynamic models feasible. However, for many questions, single-season models parameterized with appropriate fixed and random temporal effects should suffice.

Processes which generate unstructured occurrence data are complex and it is likely that the fraction of community sampling events (vs. targeted sampling) varies across eras and through time. This may be especially relevant if researchers are interested in utilizing combined museum specimen and community science records for inference. Individual collector behaviors may be important in models using these combined data-sets given that collector behavior has a potentially large effect on community science outcomes (Di Cecco et al., 2021). Incorporating these processes is an increasingly important task, as community science records far outpace the digitization and collection of specimens via other methods in the last decade (Spear et al., 2017; Shirey et al., 2021). While the current paper presents context to certain technical aspects of occupancy models that may have important consequences, there are many further complications that require additional attention. For example, restricting species’ range in multispecies occupancy modeling is a good general practice when estimating occupancy and detection. However, if researchers are interested in shifts in species ranges (e.g., range expansion), the set of modeled sites for a given species needs to be sufficiently large to allow for such range expansions. Adding a buffer around a species’ inferred range is a possible solution, but such decisions should be explored via simulation in more detail.

## Conclusion

Our analyses of simulated data demonstrate that for some unstructured data, robust estimates of temporal trends in occupancy and/or detection can be obtained, particularly when (a) community sampling events comprise at least half of all sampling events and (b) when analyses span larger temporal eras. We strongly encourage researchers to ”get to know” their data by assessing the ways in which it has been historically collected and curated and to perform, at a minimum, the guidelines we propose in this paper. Estimating the fraction of collection events that were community sampling events is critical for assessing how reliable occupancy-detection estimates may be. Overall, understanding when occupancy models are likely to fail and lead to incorrect inferences is a powerful tool when applying these models to historical museum records.

## Acknowledgements

We acknowledge funding from Simon Fraser University (to LKM and LMG), the Natural Sciences and Engineering Research Council of Canada (NSERC) (Discovery Grant to LKM), Liber Ero Fellowship Program (LMG), Georgetown University (VS), National Science Foundation Graduate Research Fellowship (VS, grant number 1937959), and the Swiss National Science Foundation fellowship (RK, grant number P2ZHP2-175028). This research was enabled in part by support from West Grid (www.westgrid.ca) and Compute Canada (www.computecanada.ca).

## Data Accessibility

The code used to generate simulated data is available via GitHub at https://github.com/lmguzman/occ_historical. A snapshot of the final code used at the time of publication is located in a Zenodo repository at xx.

## Supplementary Information

### Occupancy Model

We assume that the probability that species *i* is detected at site *j* in era *k, x_ijk_*, is drawn from a Bernoulli distribution (0 or 1) with probability (*y_ijk_*),

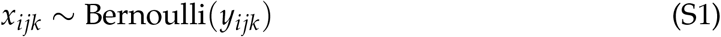

where *y_ijk_* is the product of detection probability (*p_ijk_*) and the unknown, but true occupancy state, *z_ijk_*,

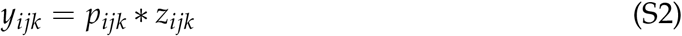

The true but unknown site occupancy for species *i* at site *j*, *z_*ijk*_* is equal to 1 if that site is occupied and 0 if it is not. We assume that this true site occupancy is drawn from a Bernoulli distribution with mean equal to the species’ occupancy probability at that site,

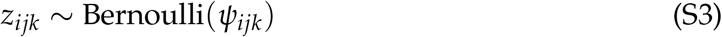

Both occupancy probability, *ψ*, and detection probability, *p*, are formulated as functions time.

We model occupancy as:

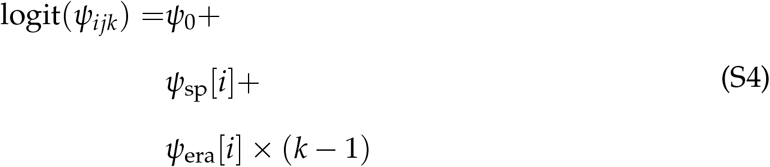

where *ψ*_0_ denotes the baseline occupancy (on the linear scale), *ψ*_sp_ [*i*] is a random species-specific intercept, and *ψ*_era_[*i*] is a random species-specific effect of era (a positive value of *ψ*_era_[*i*] would indicate that species *i* is increasing in occupancy through time). We assume that *ψ*_sp_ [*i*] and *ψ*_era_ [*i*] are normally distributed, such that:

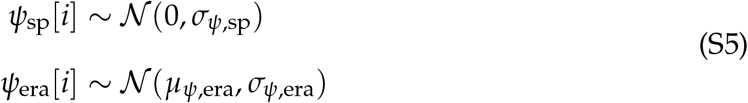

Then, we model detection probability as:

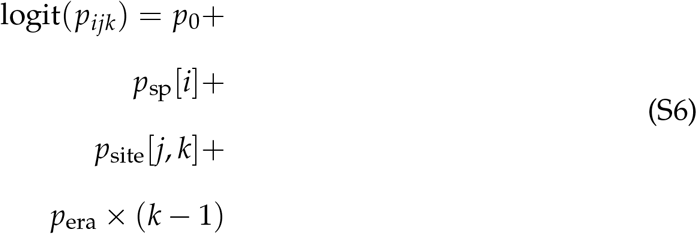

where *p*_0_ is the baseline detection probability (on the linear scale), *p*_sp_ [*i*] is a random species-specific intercept, *p*_site_[*j*, *k*] is a random site-specific intercept that varies by era, and *p*_era_ is an overall effect of era. *p*_site_ [*j*, *k*] allows spatiotemporal variability in detection probability (changing across sites and eras), and helps account for the variation that is inherent in sample effort across space and time in opportunistic historical data-sets. *p*_era_ allows detection to change systematically through time, as has likely occurred in many groups as sampling techniques have improved, for example. *p*_sp_ and *p*_site_ are assumed to be normally distributed, such that:

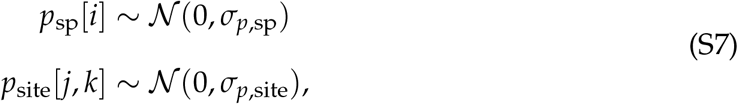

We used uninformative priors for all parameter values with normal distributions for *ψ*_0_, *μ*_*ψ*,era_, *p*_0_, *p*_era_, and uniform distributions for *σ*_*ψ*,sp_, *σ*_*ψ*,era_, *σ*_*p*,sp_, *σ*_p,site_.

We ran all models for 100,000 iterations, with a burn in period of 1000 iterations, after burn in iterations were thinned every 100 iterations, and we ran 3 chains of the MCMC. This ensured that across all of our simulations, most parameters converged (r-hat < 1.1), and this convergence was independent of parameter space and model choice.

**Table S1:** All of the parameter values used in the simulations of occurrence data.

| Parameters varied in the simulations | Values |
| --- | --- |
| Number of eras $K$ | 2, 5, 10 |
| Probability of community visits $\rho_{\text{com}}$ | 0, 0.25, 0.5, 0.75, 1 |
| Visitation probability $\mu_{v,\text{era}} / \eta_{\text{era}}$ | -0.1, 0, 0.1 |
| Species ranges | no species ranges (all species can be found in all sites), species ranges |
| Parameters values used in the simulations | Values |
| Number of species | 50 |
| Number of sites | 100 |
| Number of visits per era | 3 |
| **Detection parameters** |  |
| $p_0$ | -0.5 |
| $\sigma_{p,\text{site}}$ | 1.5 |
| $\sigma_{p,\text{sp}}$ | 0.5 |
| $p_{\text{era}}$ | 0 |
| **Occupancy parameters** |  |
| $\psi_0$ | 0 |
| $\sigma_{\psi,\text{sp}}$ | 0.5 |
| $\sigma_{\psi,\text{era}}$ | 0.2 |
| $\mu_{\psi,\text{era}}$ | 0 |
| **Visitation parameters** |  |
| $\mu_{v,0}, \eta_0$ | 0 |

### Supplementary Results

**Figure S1:**
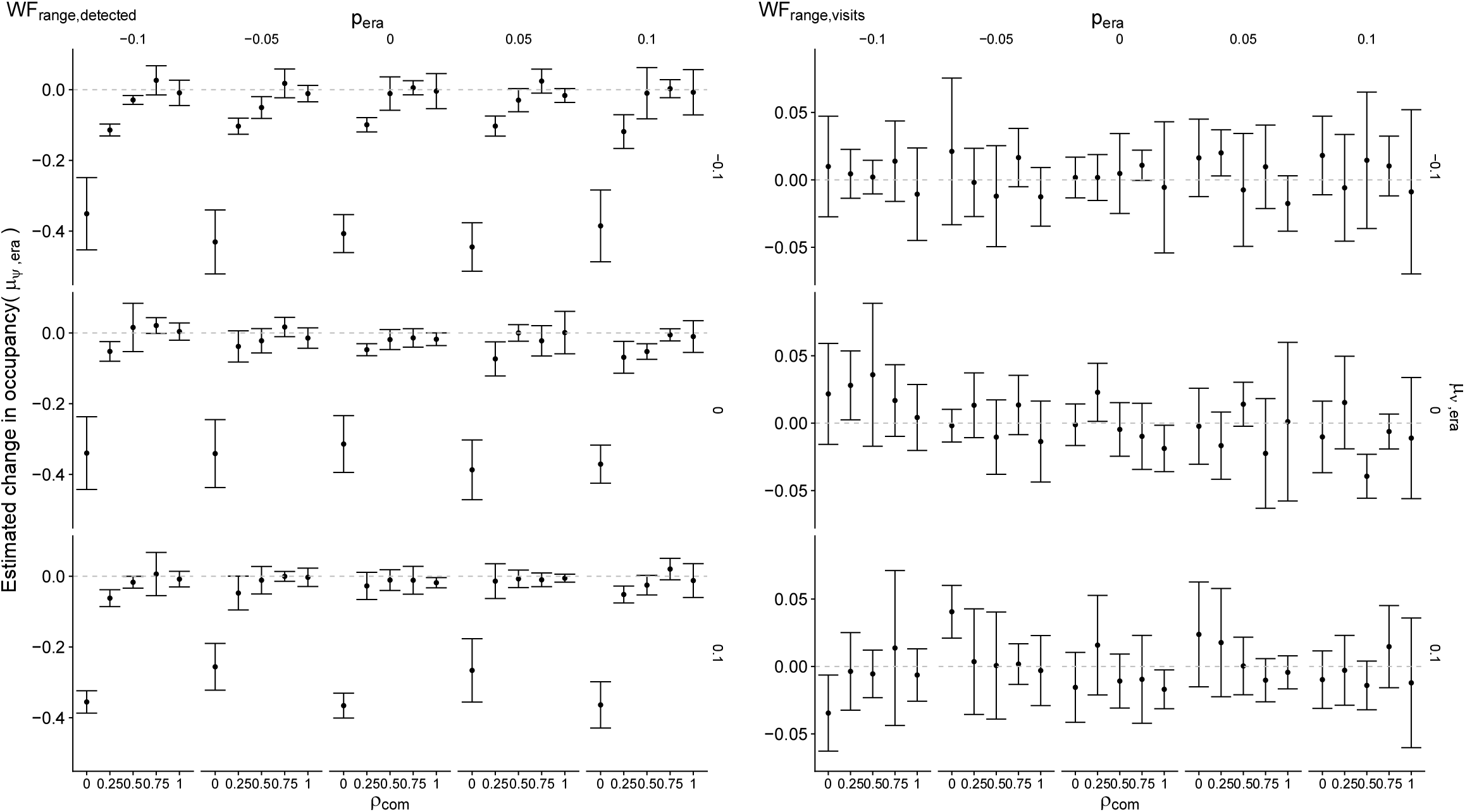
Estimated change of occupancy through time *μ*_*ψ*,era_, as a function of changes in detection through time (*p*_era_), visit history through time (*μ*_*v*,era_), and the probability of community visits (*ρ*_com_). Points are means across five replicates and error bars are the standard deviation across the replicates. The true value of *μ*_*ψ*,era_ is represented by the dashed grey line and is equal to zero.

**Figure S2:**
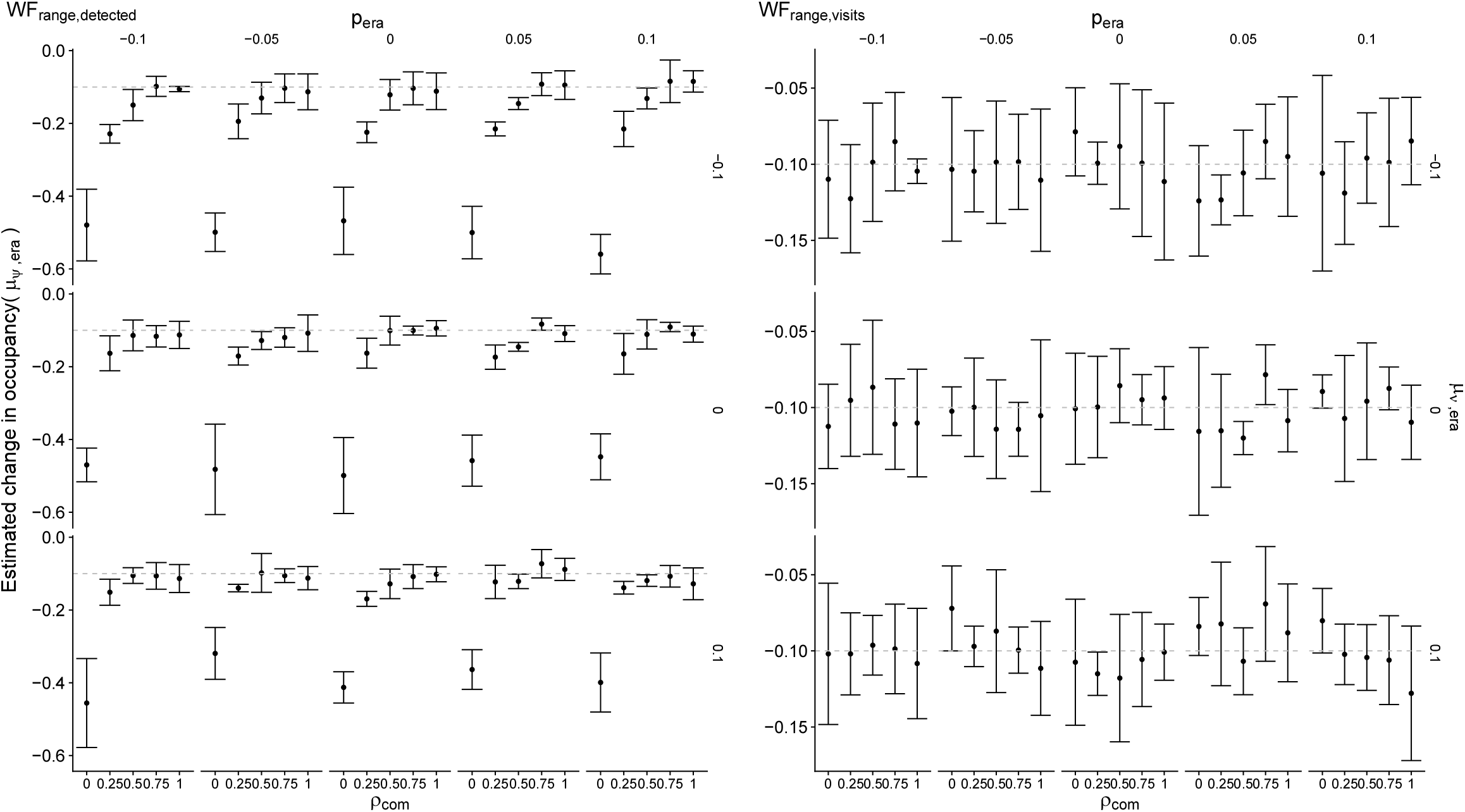
Estimated change of occupancy through time *μ*_*ψ*,era_, as a function of changes in detection through time (*p*_era_), visit history through time (*μ*_*v*,era_), and the probability of community visits (*ρ*_com_). Points are means across five replicates and error bars are the standard deviation across the replicates. The true value of *μ*_*ψ*,era_ is represented by the dashed grey line and is equal to −0.1.

**Figure S3:**
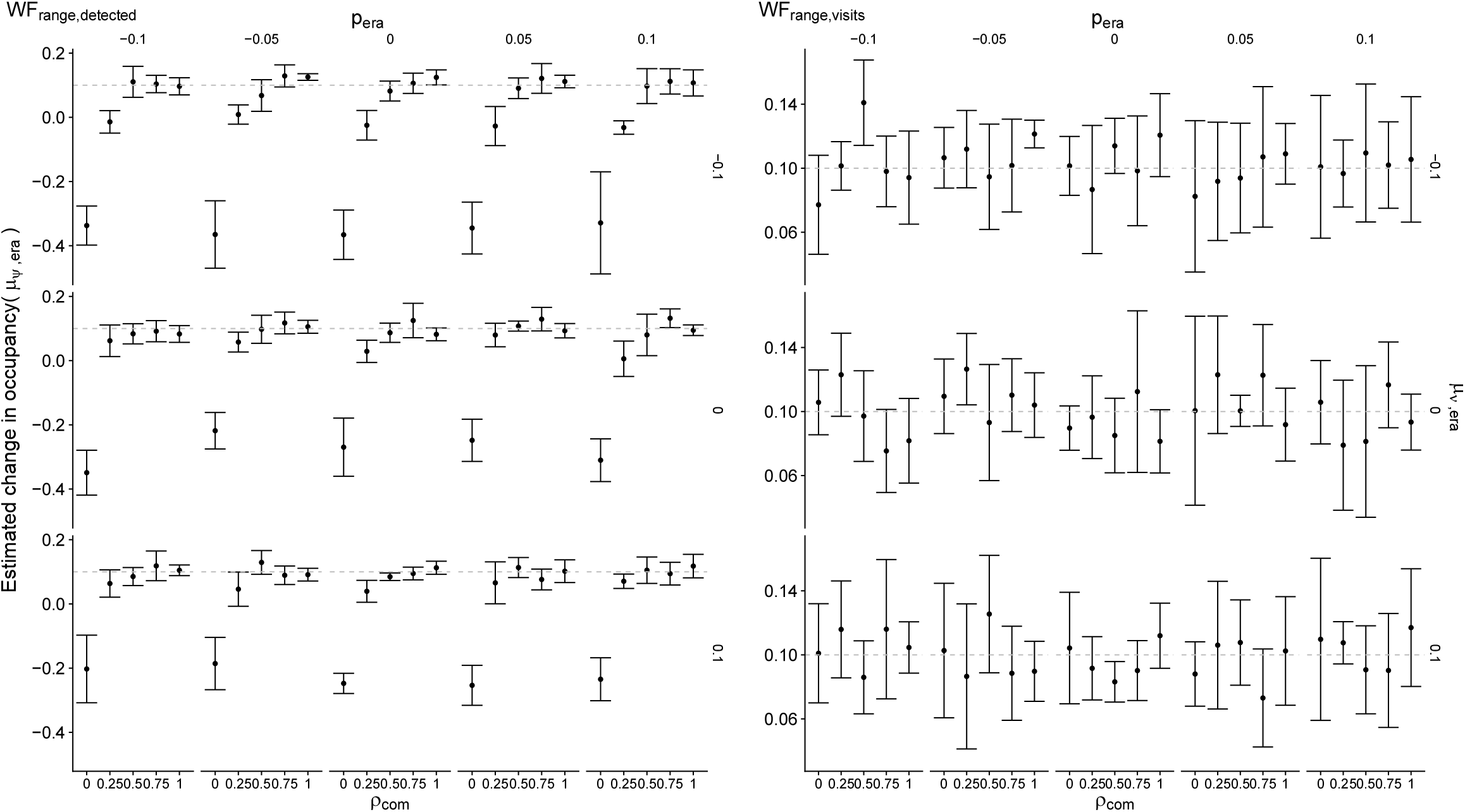
Estimated change of occupancy through time *μ*_*ψ*,era_ as a function of changes in detection through time (*p*_era_), visit history through time (*μ*_*v*,era_), and the probability of community visits (*ρ*_com_). Points are means across five replicates and error bars are the standard deviation across the replicates. The true value of *μ*_*ψ*,era_ is represented by the dashed grey line and is equal to 0.1.

**Figure S4:**
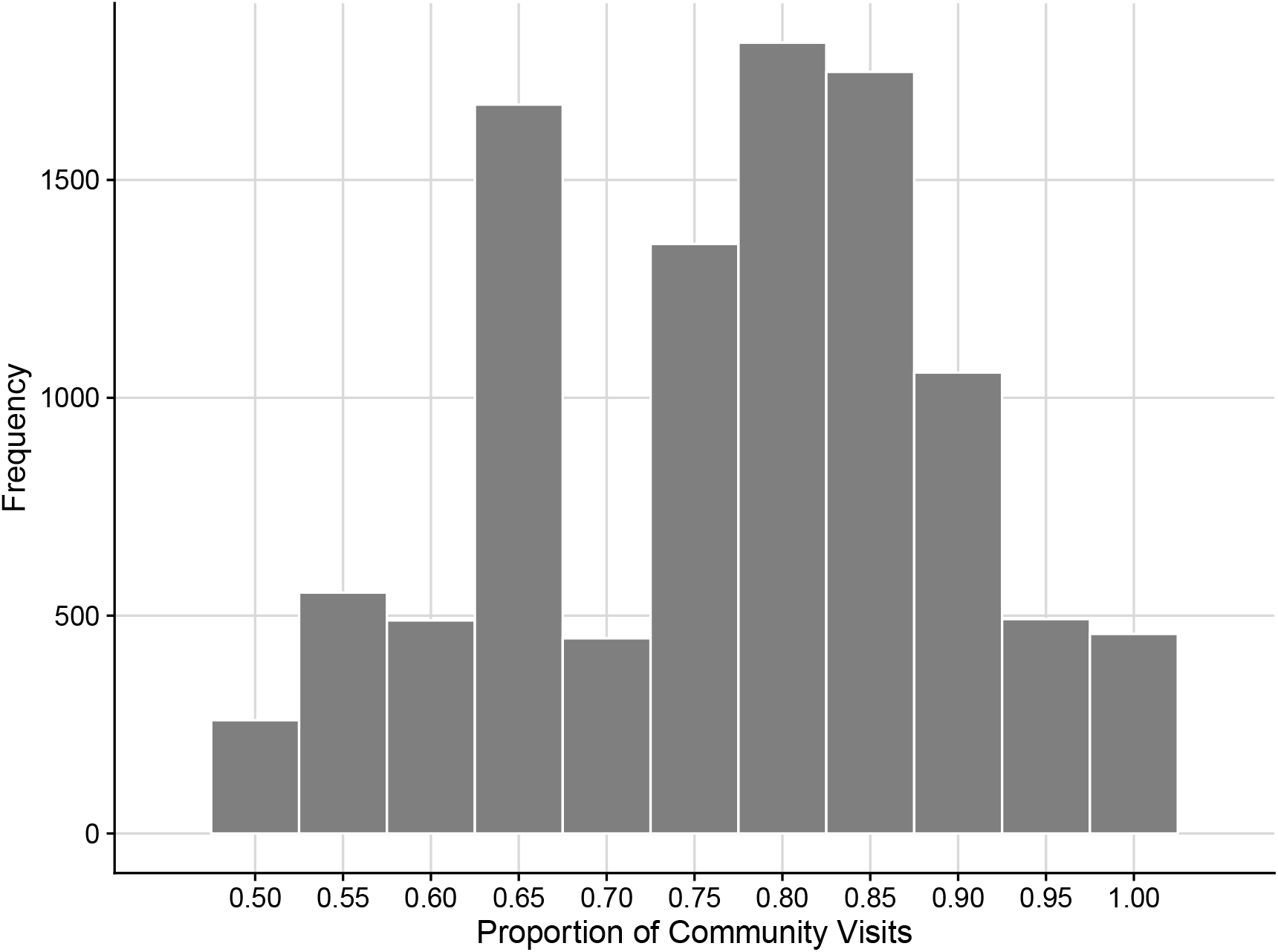
Distribution of the proportion of community visits by site/era/time interval combinations for a real-world odonate data-set. Only site/era/time interval with over half of visits as ”community sampling events” (greater than 2 species detected based on a spatiotemporal clustering algorithm) are shown.

## Notes

### Competing Interest Statement

The authors have declared no competing interest.

